# Four-dimensional tractography animates neural propagations via distinct interhemispheric pathways

**DOI:** 10.1101/2020.12.11.421495

**Authors:** Takumi Mitsuhashi, Masaki Sonoda, Jeong-won Jeong, Brian H. Silverstein, Hirotaka Iwaki, Aimee F. Luat, Sandeep Sood, Eishi Asano

## Abstract

**Objective:** To visualize and validate the dynamics of interhemispheric neural propagations induced by single-pulse electrical stimulation (SPES).

**Methods:** This methodological study included three patients with drug-resistant focal epilepsy who underwent measurement of cortico-cortical spectral responses (CCSRs) during bilateral stereo-electroencephalography recording. We delivered SPES to 83 electrode pairs and analyzed CCSRs recorded at 268 nonepileptic electrode sites. Diffusion-weighted imaging (DWI) tractography localized the interhemispheric white matter pathways as streamlines directly connecting two electrode sites. We localized and visualized the putative SPES-related fiber activation, at each 1-ms time window, based on the propagation velocity defined as the DWI-based streamline length divided by the early CCSR peak latency.

**Results:** The resulting movie, herein referred to as four-dimensional tractography, delineated the spatiotemporal dynamics of fiber activation via the corpus callosum and anterior commissure. Longer streamline length was associated with delayed peak latency and smaller amplitude of CCSRs. The cortical regions adjacent to each fiber activation site indeed exhibited CCSRs at the same time window.

**Conclusions:** Our four-dimensional tractography successfully animated neural propagations via distinct interhemispheric pathways.

**Significance:** Our novel animation method has the potential to help investigators in addressing the mechanistic significance of the interhemispheric network dynamics supporting physiological function.

## 1. Introduction

The goal of this methodological study is to generate an animation movie that accurately visualizes the rapid dynamics of interhemispheric neural propagations in the human brain. Studies using diffusion-weighted imaging (DWI) tractography suggest that the healthy brain includes the interhemispheric white matter pathway directly connecting two hemispheres, including the corpus callosum and anterior commissure (Catani *et al*., 2002; Hofer and Frahm, 2006; McNab *et al*., 2013). Invasive studies of epilepsy patients have reported that single-pulse electrical stimulation (SPES) of the nonepileptic cerebral cortex can elicit neural responses at contralateral homotopic regions within 50 ms (Wilson *et al*., 1990; 1991; Umeoka *et al*., 2009; Terada *et al*., 2012; Mălîia *et al*., 2018; Trebaul *et al*., 2018; Dionisio *et al*., 2019). In other words, these studies indicate that single-axonal neural propagations can be completed in 50 ms in the majority of the interhemispheric white matter pathways. Collective evidence suggests that interhemispheric neural networks support variable sensorimotor and cognitive function (Funnell *et al*., 2000; Catani *et al*., 2002; Aralasmak *et al*., 2006; Andoh *et al*., 2015; Roland *et al*., 2017; Rossini *et al*., 2019). Conversely, interhemispheric propagation of epileptic discharges is responsible, at least in part, for defining the seizure semiology as well as the worsening of cognitive impairment in patients with epilepsy (Gotman, 1987; Lieb *et al*., 1991; Adam *et al*., 1994; Ono *et al*., 2011; Peltola *et al*., 2011; Unterberger *et al*., 2016). In summary, we expect that our study will help investigators in addressing the mechanistic relevance of the interhemispheric network dynamics in the physiological function and the propagation of epileptic discharges.

In the present study of epilepsy patients who underwent bilateral stereoelectroencephalography (sEEG) recording, we localized the interhemispheric white matter pathways as streamlines directly connecting two electrode sites using DWI tractography (Trebaul *et al*., 2018; Yeh *et al*., 2018). We delivered SPES to pairs of electrode contacts and measured cortico-cortical spectral responses (CCSRs; Sugiura *et al*., 2020; Mitsuhashi *et al*., 2020) in distant sites, including those in the contralateral hemisphere. The early CCSR peak taking place within 11-50 ms post-stimulus is suggested to reflect a neural response induced by single-axonal propagation from a given SPES site (Matsumoto *et al*., 2017; Usami *et al*., 2019; Silverstein *et al*., 2020). A high magnitude of CCSR reflects the presence of effective connectivity via which SPES sites can transfer large neural signals directly to given recording sites (Sugiura *et al*., 2020). The present study localized and animated the putative SPES-related fiber activation based on the propagation velocity defined as the DWI-based streamline length divided by the early CCSR peak latency, as performed in our previous study of *intrahemispheric* neural propagations (Silverstein *et al*., 2020). We referred to the resulting animation movie as four-dimensional (4D) tractography. We addressed the following hypotheses to test the validity of our 4D tractography data. [1] We hypothesized that the corpus callosum would be found to connect the SPES and CCSR sites within the frontal lobes, whereas the anterior commissure would connect those within the temporal lobes. Tractography studies suggest that substantial proportions of the frontal lobes are connected interhemispherically via the anterior-middle portions of the corpus callosum, whereas the temporal lobes via the anterior commissure (Catani *et al*., 2002; Aralasmak *et al*., 2006). [2] We hypothesized that a longer DWI-based streamline would be associated with delayed CCSR peak latency and smaller CCSR amplitude. [3] We hypothesized that the cortical contacts adjacent to each putative fiber activation site on the 4D tractography would exhibit CCSRs at the same time window.

## 2. Methods

### 2.1. Participants

We studied the same cohort of three patients with drug-resistant focal epilepsy who underwent measurement of CCSRs during bilateral sEEG recording (age range: 12-16 years; **Table 1**; Mitsuhashi *et al*., 2020). The Institutional Review Board at Wayne State University has approved the present study. We obtained informed consent from the legal guardians of patients and assent from patients.

**Table 1.**
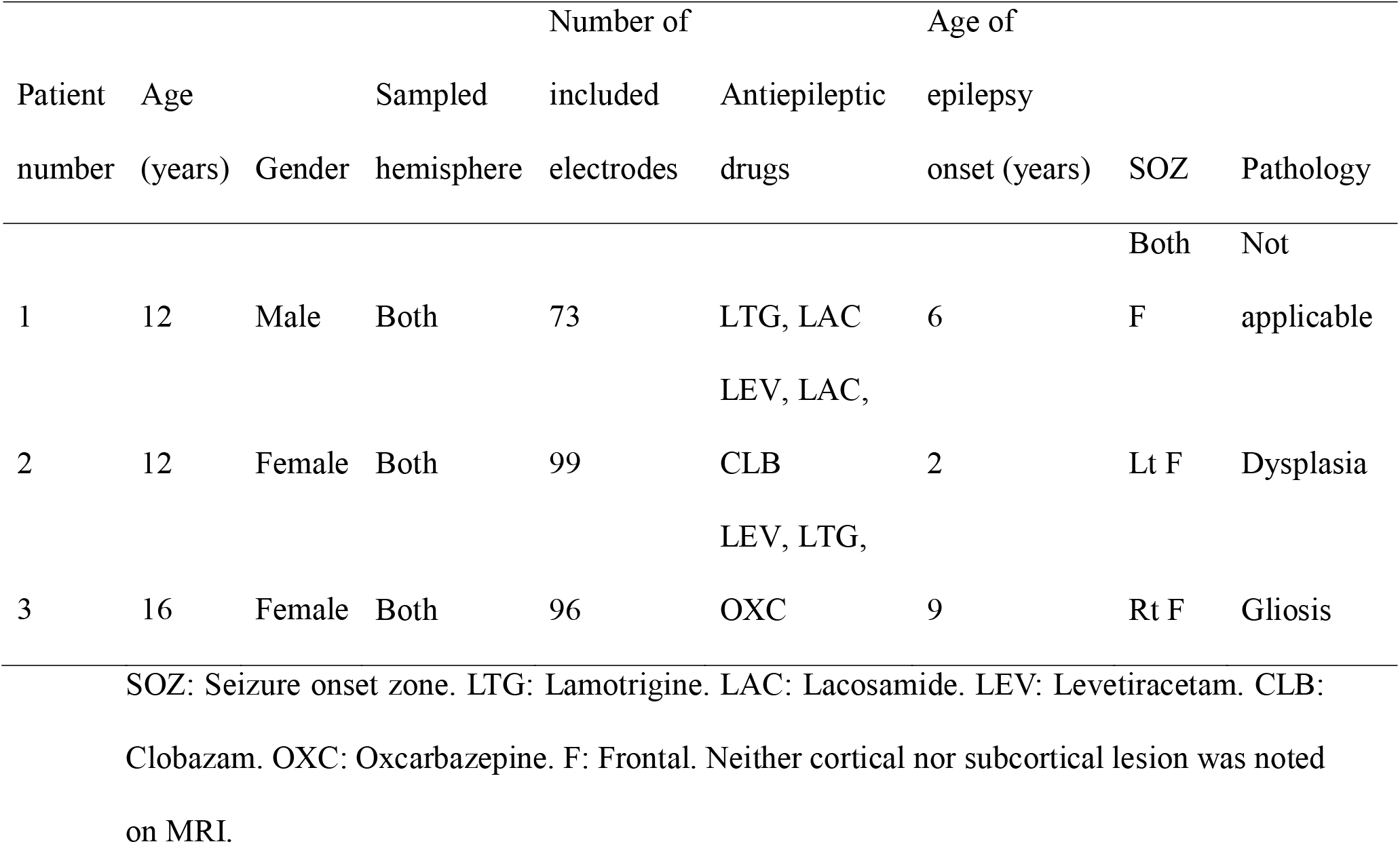
Patients profile.

### 2.2. Intracranial electrode placement

All patients had platinum depth electrodes implanted stereotactically under MRI guidance in both hemispheres. Each depth electrode had eight, 12, and 16 contacts (length: 2 mm; center-to-center distance: 3.5, 3.5, and 4.43 mm, respectively; PMT, Chanhassen, MN, USA).

### 2.3. Imaging process

Using a pre-implant T1-weighted spoiled gradient-recalled echo sequence MRI and a post-implant CT image, we created a fusion image accurately denoting sEEG depth electrodes within the brain (Mitsuhashi *et al*., 2020). We used the Statistical Parametric Mapping 12 (http://www.fil.ion.ucl.ac.uk/spm/software/spm12/) to spatially normalize each electrode to the standard Montreal Neurological Institute (MNI) space. We determined the anatomical location of each depth electrode based on the automated anatomical labeling atlas (Tzourio-Mazoyer *et al*., 2002) implemented in the FieldTrip toolbox (http://www.fieldtriptoolbox.org) (**Fig. 1; Video S1**).

**Fig. 1.**
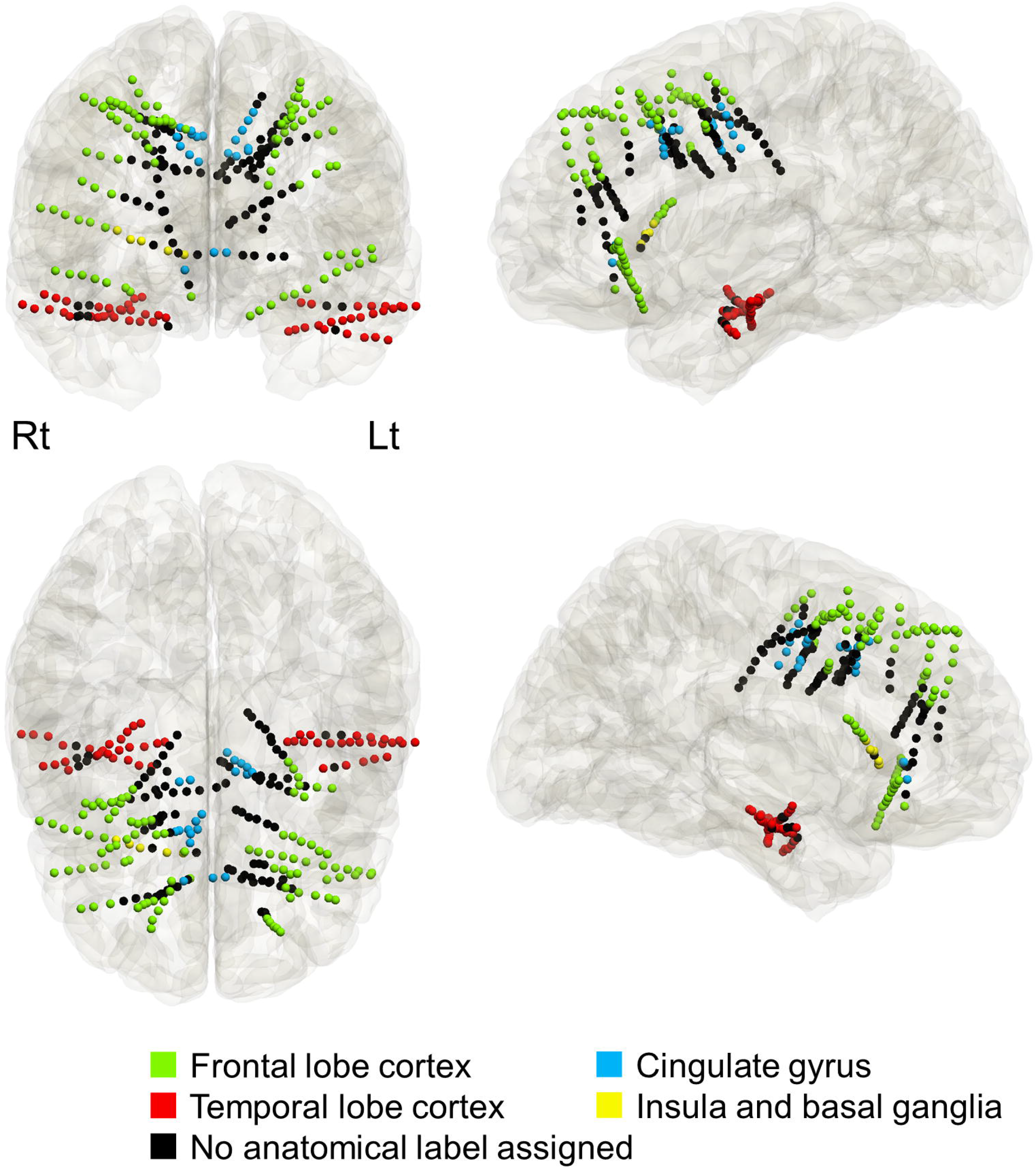
The anatomical locations of depth electrode contacts included in the analysis. Light green: frontal lobe cortex. Light blue: cingulate gyrus. Red: temporal lobe cortex. Yellow: insula and basal ganglia. Black: no anatomical label assigned (i.e., white matter).

### 2.4. Extraoperative sEEG recording

We acquired sEEG data using the method identical to that reported previously (Mitsuhashi *et al*., 2020). The sEEG sampling rate was 1,000 Hz, and the bandpass was 0.016 - 300 Hz. Visual assessment of sEEG signals was performed on bipolar montage. The present study excluded the seizure onset zone (SOZ; Asano *et al*., 2009), irritative zone (Kural *et al.,* 2020), electrode contacts outside the brain, and artifactual contacts from further analyses. Previous SPES studies reported that stimulation at the SOZ elicited accentuated cortico-cortical evoked potentials (CCEPs; Iwasaki *et al*., 2010; Enatsu *et al*., 2012).

### 2.5. Single-pulse electrical stimulation (SPES)

We performed SPES using the method previously reported (Mitsuhashi *et al*., 2020). In each SPES trial, we delivered a train of electrical stimuli to an adjacent pair of sEEG electrodes at a frequency of 1 Hz for 40 s during a resting period of sleep. Each stimulus consisted of a square wave pulse of 0.3 ms duration, 5 mA intensity, and biphasic polarity. No adverse events related to the SPES procedures were observed or reported.

### 2.6. Measurement of early CCSR

We quantified the magnitude of CCSR_20-60 Hz_ using a time-frequency analysis similar to those reported previously (Tallon-Baudry *et al*., 1997; Oostenveld *et al*., 2011; Mitsuhashi *et al*., 2020). We excluded recording contacts within 1 cm from the SPES site from analysis (Sugiura *et al*., 2020; Prime *et al*., 2020). The Morlet wavelet method, as implemented in the FieldTrip toolbox, transformed sEEG signals into time-frequency bins (2 Hz frequency bins; three cycles for each frequency; sliding in 1 ms steps) within a period of 11-50 ms with a bandpass of 20-60 Hz. We computed the percent change of CCSR amplitude relative to that during the 50-200 ms pre-stimulus baseline period for each frequency bin and averaged across 40 stimuli. CCSRs satisfying the following three-step criteria were considered to be significant (Mitsuhashi *et al*., 2020). [Reproducibility] The CCSR amplitude augmentation must be reproducible based on the cluster-based permutation test (Maris and Oostenveld, 2007). [Magnitude of augmentation] The CCSR amplitude must be augmented at least by 80% compared to the baseline period. [Exclusion of far-field signals] The CCSR amplitude on bipolar montage must be not smaller than 30% of that on common average montage. Our recent sEEG study revealed that extra-brain contacts occasionally showed CCSRs visible on common average but not on bipolar montage (Mitsuhashi *et al*., 2020). We finally computed the early CCSR peak latency using the exact timing showing the maximum amplitude augmentation within the 11-50 ms window (Sugiura *et al*., 2020).

### 2.7. Diffusion-weighted imaging (DWI) tractography

In this study of CCSR-based effective connectivity between nonepileptic regions, we generated DWI tractography using the data averaged across the 1,065 healthy individuals participating in the Human Connectome Project (http://brain.labsolver.org/diffusion-mri-templates/hcp-842-hcp-1021; Yeh *et al*., 2018). We placed seed regions (5-mm radius spheres) at the midpoint of given SPES pairs and a recording contact showing a significant early CCSR. We used DSI Studio (http://dsi-studio.labsolver.org/) to generate streamlines directly connecting each pair of seed regions within the common MNI space (http://www.bic.mni.mcgill.ca/ServicesAtlases/ICBM152NLin2009). Streamlines satisfying the following criteria were considered to be significant: a fractional anisotropy threshold of 0.2, a maximum turning angle of 70°, step size of 0.3 mm, and streamline length from 10 to 250 mm. The shortest significant streamline connecting each pair of seed regions was used for further analysis. We classified a given streamline into one of the following nine commissural pathways based on the MNI coordinate of the streamline in the median plane: [1] rostrum, [2] genu, [3] rostral body, [4] anterior midbody, [5] posterior midbody, [6] isthmus, [7] splenium of the corpus callosum, [8] anterior commissure, and [9] hippocampal commissure (**Fig. 2;** Witelson, 1989; Fonov *et al*., 2011).

**Fig. 2.**
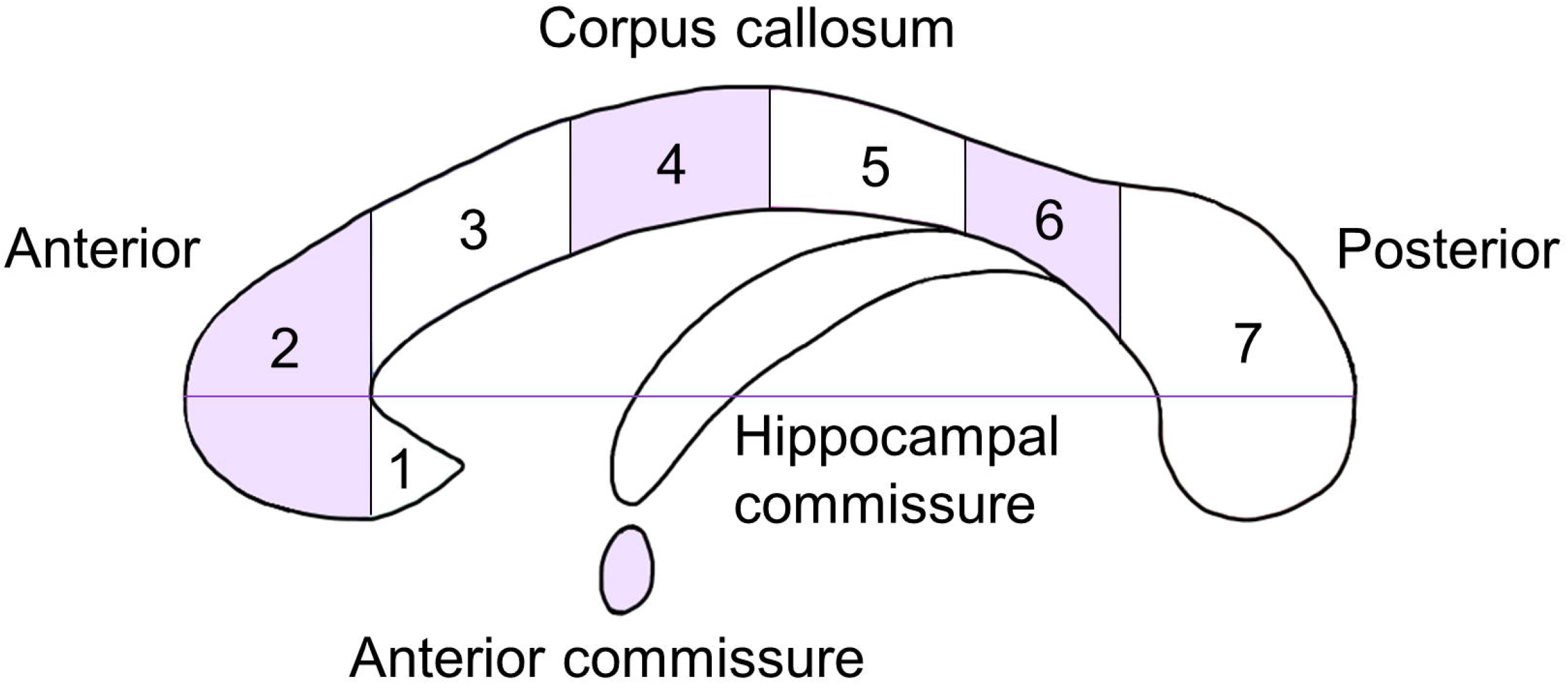
Commissural pathways. The corpus callosum consisted of [1] rostrum, [2] genu, [3] rostral body, [4] anterior midbody, [5] posterior midbody, [6] isthmus, and [7] splenium.

### 2.8. 4D tractography

We animated the spatiotemporal dynamics of SPES-related neural propagations on the DWI-based streamlines in the MNI space. We defined the propagation velocity as the entire streamline length divided by the early CCSR peak latency (Silverstein *et al*., 2020). We defined the putative fiber activation as the white matter site showing neural propagation at a given moment (**Fig. 3**). The velocity measure effectively allowed us to localize the putative fiber activation at each 1-ms time window and to visualize the temporal changes of fiber activation sites along the streamline.

**Fig. 3.**
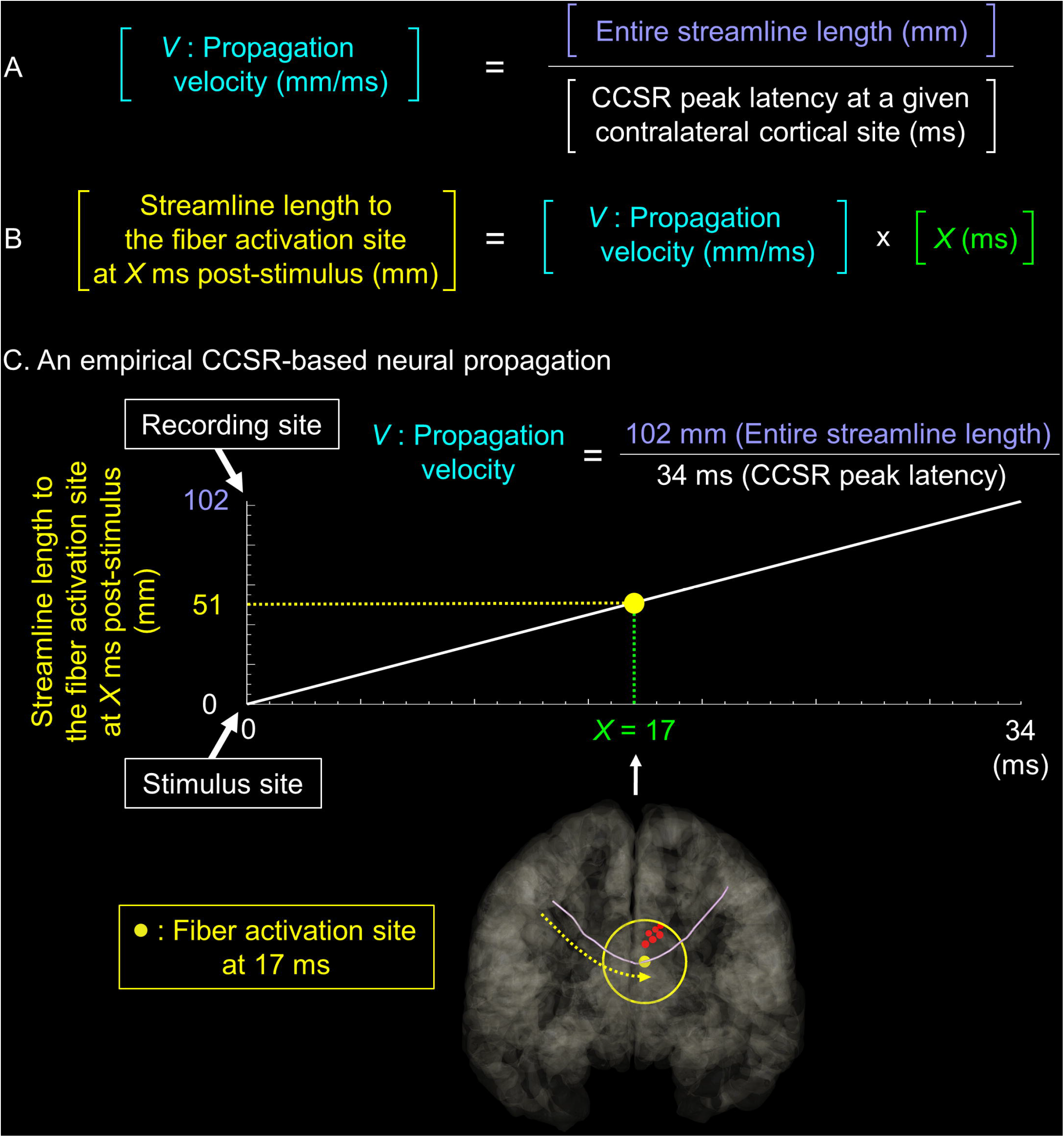
Definition of putative fiber activation site. The velocity (*V*) of single-pulse electrical stimulation (SPES)-related neural propagation was defined as the entire streamline length divided by the peak latency of cortico-cortical spectral response (CCSR) recorded at a contralateral cortical site receiving a given streamline. (B) A fiber activation site at *X* ms post-stimulus was localized within the streamline *V* * *X* mm away from the stimulus site. (C) An empirical CCSR-based neural propagation via the corpus callosum is presented. The velocity of SPES-related neural propagation was 3.0 mm/ms (i.e., 102 mm / 34 ms). The yellow dot denotes the fiber activation site at 17 ms post-stimulus (i.e., 3.0 * 17 mm away from the stimulus site). Red dots: cortical contacts within 2 cm from the fiber activation site at 17 ms.

### 2.9. Association between the DWI-based streamline length and CCSR measures

A mixed model analysis tested the hypothesis that longer streamline length would be associated with delayed CCSR peak latency as well as smaller CCSR amplitude. The dependent variables included the early CCSR peak latency and CCSR amplitude. The fixed effect predictor variable was DWI-based streamline length. The random factors included patients and intercept. Statistical analyses were performed using IBM SPSS Statistics version 22 (IBM Corp., Armonk, NY, USA). The significance was set at a p-value of 0.05.

### 2.10. Assessment of the CCSRs at cortical regions adjacent to a given fiber activation

We validated our 4D tractography method to localize fiber activations based on the velocity of SPES-related neural propagation. The mixed model analysis determined whether the peak latency of CCSRs at cortical regions adjacent to each fiber activation site would be predicted by the latency of neural propagation reaching to a given white matter site. We calculated the CCSR size at a given moment, averaged across cortical contacts within 2 cm from a given fiber activation site (**Fig. 3**). Thereby, the CCSR size was z-score transformed across 11-to 50-ms post-stimulus periods. The dependent variable was the peak latency of CCSRs at cortical contacts adjacent to a fiber activation site. The fixed effect predictor variable was the latency of propagation to a fiber activation site. The random factors included patients and intercept. This analysis effectively determined whether CCSRs would be indeed augmented at surrounding cortical contacts when SPES-related neural signals are propagated to a white matter site.

### 2.11. Data and code availability

All sEEG data, as well as the Matlab-based code used to generate the 4D tractography, are available upon request to the corresponding author.

## 3. Results

### 3.1. Configurations and anatomical locations of implanted electrodes

Ninety-six electrode contacts were classified to be in the frontal lobe cortex, 21 in the cingulate gyrus, 52 in the temporal lobe cortex, five in the insula/basal ganglia (**Fig. 1; Videos S1**). The remaining 94 contacts were within the white matter. SPES was delivered to 83 electrode contact pairs in total. The cumulative number of CCSR recording channels was 7,325.

### 3.2. Visualization of neural propagations via distinct interhemispheric streamline pathways

We localized 56 interhemispheric streamlines via which SPES elicited significant CCSRs in the contralateral hemisphere (**Fig. 4**). The 4D tractography successfully animated the dynamics of SPES-related neural propagations taking place via the anterior commissure and corpus callosum, respectively (**Video S2; Fig. 5**).

**Fig. 4.**
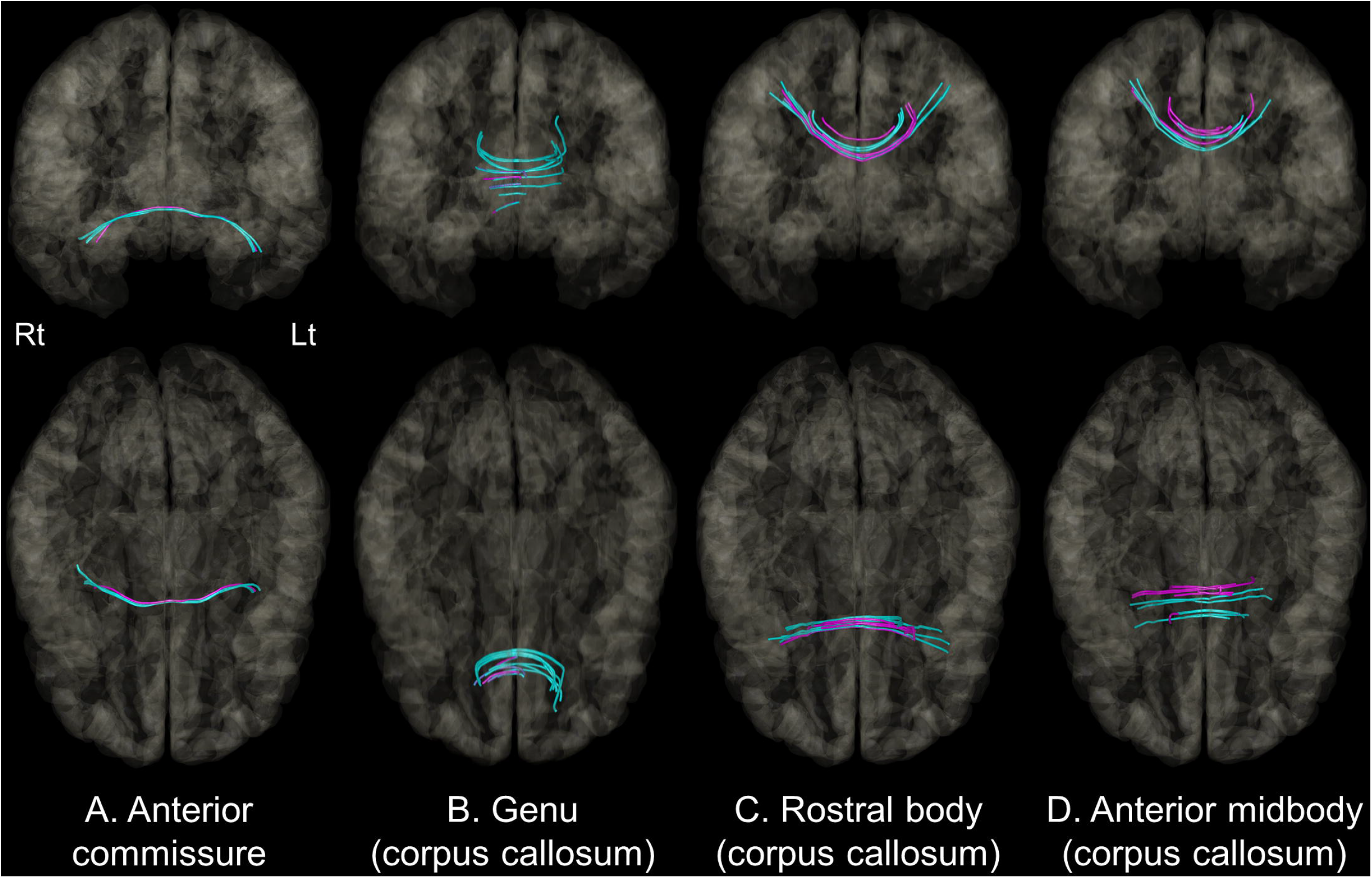
Interhemispheric white matter pathways visualized as streamlines connecting single-pulse electrical stimulation (SPES) and cortico-cortical spectral response (CCSR) sites. Interhemispheric streamline pathways via (A) the anterior commissure, (B) the genu, (C) the rostral body, and (D) the anterior midbody of the corpus callosum. Magenta: streamlines between SPES sites in the left hemisphere and CCSR sites in the right hemisphere. Cyan: streamlines between SPES in the right hemisphere and CCSR sites in the left hemisphere.

**Fig. 5.**
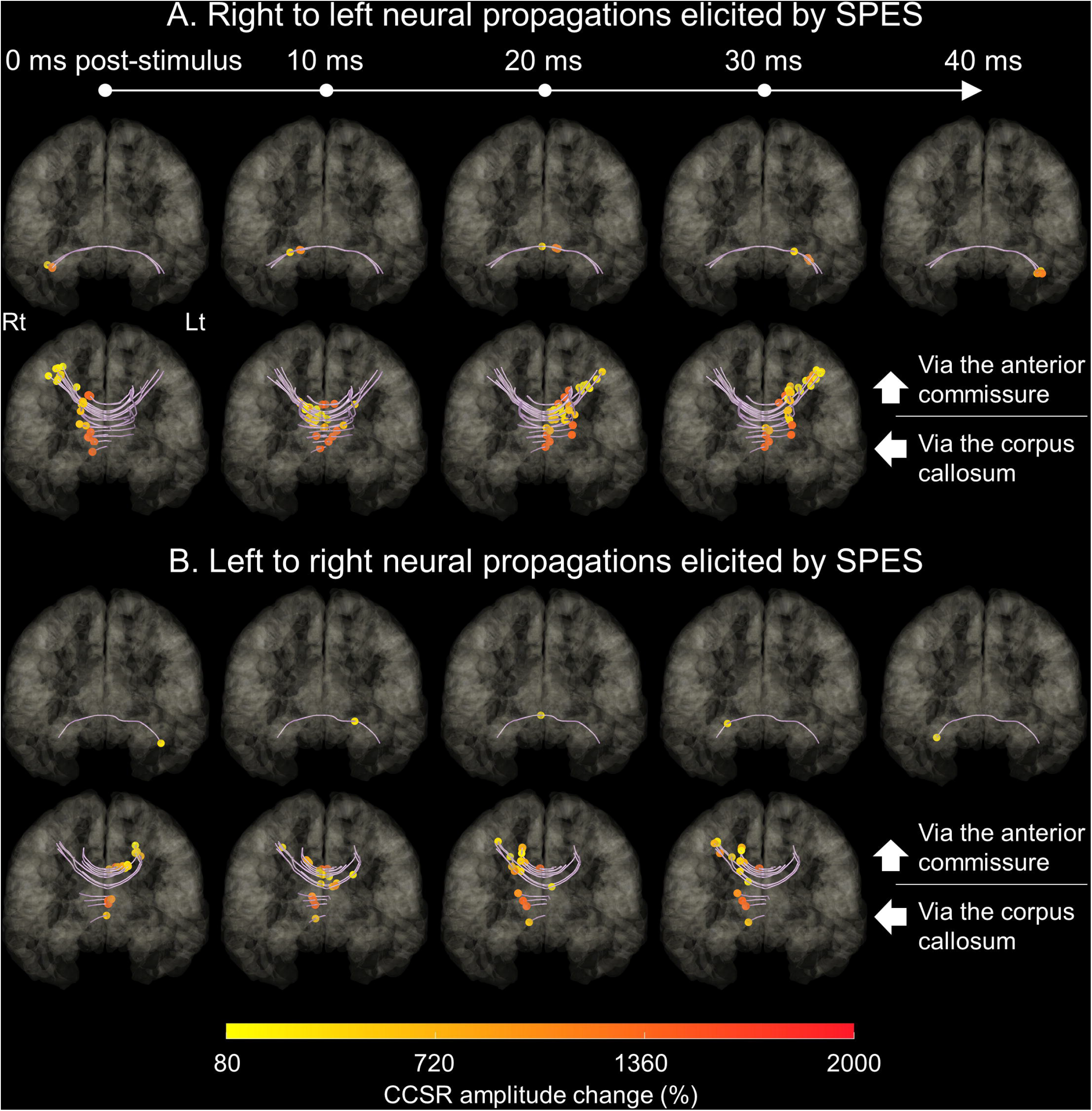
Snapshots of four-dimensional (4D) tractography. The 4D tractography, derived from the early cortico-cortical spectral response (CCSR) and diffusion-weighted imaging data, animates the dynamics of single-axonal neural propagations elicited by single-pulse electrical stimulation (SPES). (A) The snapshots of right-to-left neural propagations at 0, 10, 20, 30, and 40 ms post-stimulus. Circles: Putative fiber activation at a given moment. Circle color: CCSR amplitude change (%) measured at the recording site in the contralateral hemisphere. (B) The snapshots of left-to-right neural propagations.

Five streamlines passed through the anterior commissure; thereby, all of them connected the bilateral anterior temporal lobes (mean early CCSR peak latency: 37.2 ms [SE: 1.4]; mean propagation velocity: 2.46 mm/ms [SE: 0.05]). The remaining 51 streamlines passed through the corpus callosum, and all of them connected either bilateral frontal lobes or bilateral anterior cingulate gyri (mean early CCSR peak latency: 21.9 ms [SE: 1.0]; mean propagation velocity: 2.41 mm/ms [SE: 0.16]). Fisher’s exact probability test suggested a double dissociation between the interhemispheric white matter pathways and the origin/destination of SEPS-related neural propagations (p<0.0001). Of 51 streamlines passed through the corpus callosum, seventeen streamlines passed through the genu, 19 through the rostral body, and 15 through the anterior midbody of the corpus callosum.

### 3.3. Association between the DWI-based streamline length and CCSR measures

Longer DWI-based streamline length was associated with delayed CCSR peak latency (mixed model estimate = +0.179 ms/mm; t = +6.150; p < 0.001) and smaller CCSR amplitude (mixed model estimate = −36.2 %/mm; t = −5.784; p < 0.001; **Fig. 6**).

**Fig. 6.**
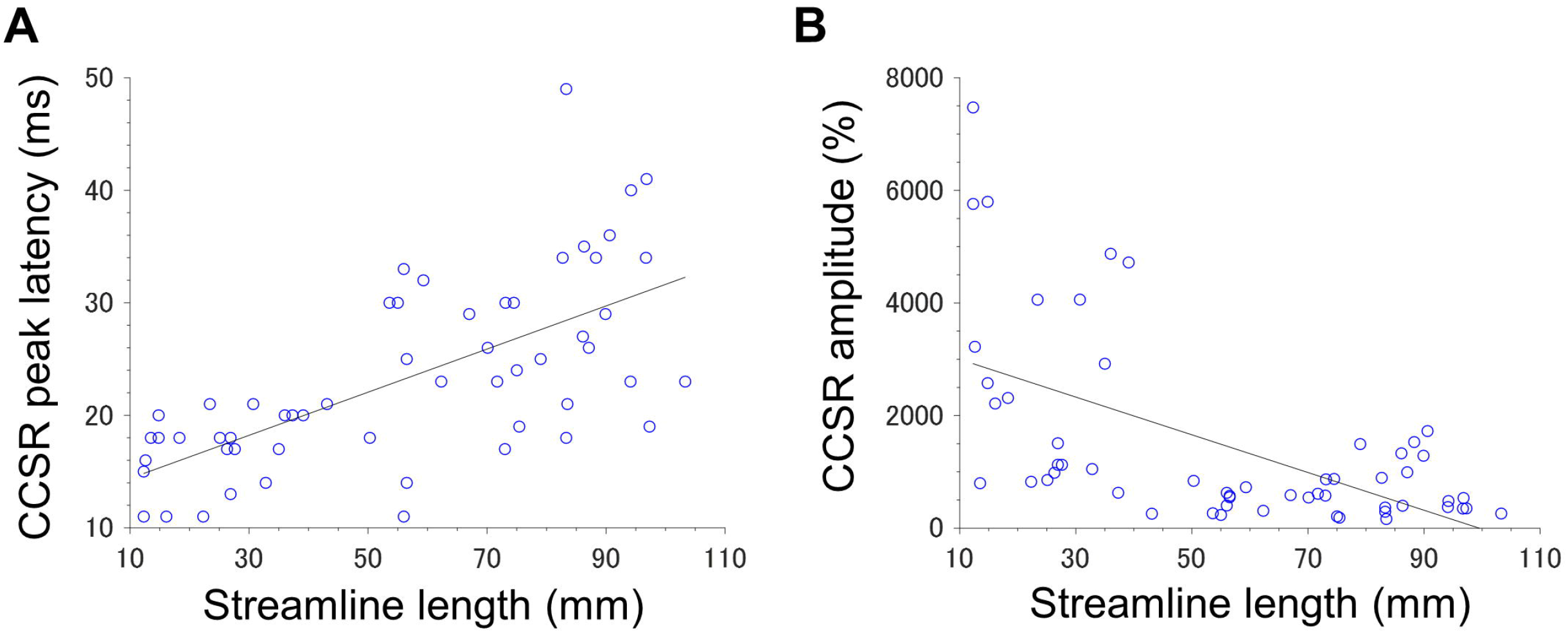
Association between the tractography-derived streamline length and cortico-cortical spectral response (CCSR) measures. (A) Each circle denotes the relationship between the streamline length and CCSR peak latency. (B) Each denotes the relationship between the streamline length and CCSR amplitude. Solid line: the best-fit to our CCSR/tractography data.

### 3.4. CCSRs at cortical regions adjacent to a fiber activation

Cortical contacts adjacent to a fiber activation showed CCSRs maximally augmented when SPES-related neural signals were propagated to the neighboring white matter site. For example, 17 streamlines had a fiber activation at 26-ms post-stimulus; thereby, cortical contacts within 2 cm from the fiber activation showed the CCSR maximally augmented exactly at 26-ms post-stimulus (**Fig. 7**). The mixed model analysis revealed that the peak latency of CCSRs at cortical contacts adjacent to a fiber activation site was predicted by the latency of SPES-related signal reaching a given fiber activation site (t = +21.404; p < 0.001).

**Fig. 7.**
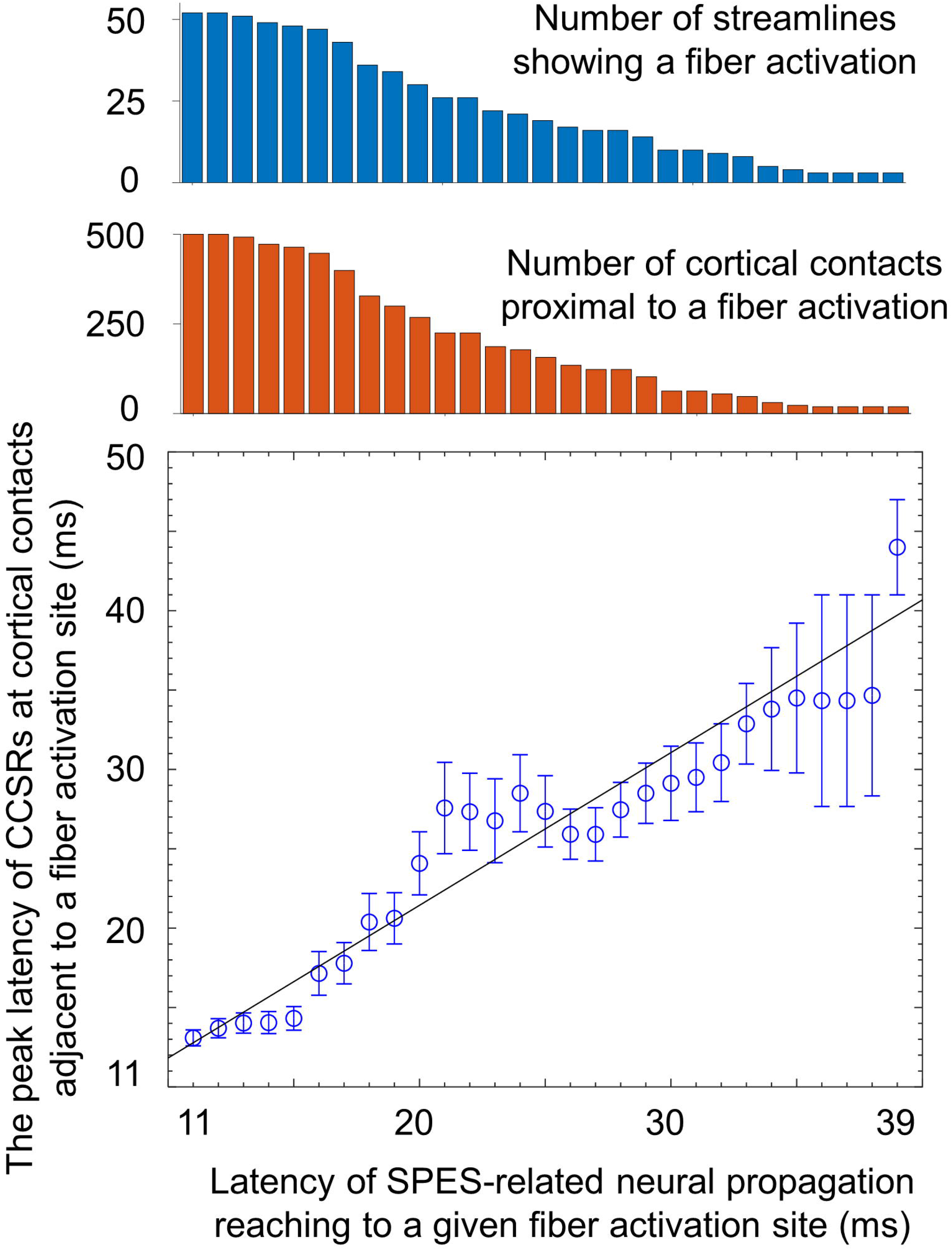
Relationship between the putative fiber activation sites and the peak latency of cortico-cortical spectral responses (CCSRs) at adjacent cortical contacts. X-axis: Latency of single-pulse electrical stimulation (SPES)-related neural propagation reaching to a given fiber activation site. We excluded fiber activations taking place at a latency of ≥40 ms since no more than two streamline pathways were available. Y-axis: The peak latency of CCSRs at cortical contacts adjacent (≤2 cm) to a fiber activation site. Fifty-two streamlines showed a putative fiber activation at 11-ms post-stimulus, whereas only three streamlines showed one at 39-ms. Solid line: the best-fit to our CCSR/tractography data based on the linear regression model (Y = 0.962X +2.2). Bar: standard error.

## 4. Discussion

### 4.1. Significance of 4D tractography in understanding the interhemispheric network dynamics

To our best knowledge, this is the first-ever study that animated the rapid dynamics of interhemispheric neural propagations taking place via the anterior commissure and corpus callosum. Our study demonstrated a double dissociation between the white matter pathways and the cortical origin/destination of neural propagations. Namely, SPES of anterior temporal lobe sites elicited CCSRs in the contralateral anterior temporal regions via the anterior commissure. Conversely, SPES of frontal lobe/anterior cingulate sites elicited CCSRs in the contralateral homotopic regions via the corpus callosum. Our results complement previous SPES studies characterizing the strength and latency of interhemispheric neural propagations between the bilateral mediobasal temporal lobes as well as the bilateral frontal lobes (Wilson *et al*., 1990; 1991; Umeoka *et al*., 2009; Terada *et al*., 2012; Trebaul *et al*., 2018). The novelty of our SPES study lies in the visualization of the temporal dynamics of fiber activations taking place within specific white matter bundles with a time resolution of 1 ms at a group level (**Video S2**). The 4D tractography might be useful for laypeople to understand how each hemisphere can communicate with each other.

Our novel animation method has the potential to help investigators in addressing the mechanistic significance of the interhemispheric network dynamics supporting physiological function. Investigators have reported that corpus callosotomy is an effective method to reduce the severity of astatic and tonic seizures, whereas the risk of postoperative neurological deficits is not negligible (Luat *et al*., 2017). Potential neurological symptoms related to callosal disconnection include apraxia of the non-dominant hand as well as impaired naming of objects presented in the non-dominant visual field or held by the non-dominant hand (Feinberg *et al*., 1992; Geschwind *et al*., 1995; Suzuki *et al*., 1998; Catani and ffytche, 2005). These investigators infer that the interhemispheric network plays an essential role in some of the sensorimotor and cognitive functions. They also suggest that information processed in the sensory cortex of the non-dominant hemisphere would be transferred to the dominant hemisphere via a commissural pathway prior to relevant motor responses. Our 4D tractography can be used to visualize the location of a fiber activation transmitted from a given site of interest, including the eloquent cortex. We hypothesize that analysis of the spatio-temporal relationship between the underlying cortical function and interhemispheric effective connectivity patterns might delineate the intra- and inter-hemispheric pathways supporting the functions necessary to execute a cognitive task. Unlike that of subdural recording, the extent of sEEG sampling is generally restricted in a horizontal dimension. Thus, multicenter studies of large datasets are needed to generate a dynamic atlas showing the interhemispheric connectivity at the whole-brain level and to address some of the outstanding questions raised by investigators.

### 4.2. Methodological considerations

We validated the location of given fiber activations delineated on the 4D tractography. As best presented in **Fig. 7**, the CCSR amplitude at cortical contacts adjacent to a given fiber activation was maximized at a given propagation latency. This validation procedure is the best we can do since the assessment of neural responses at the white matter is difficult (Mercier *et al*., 2017; Mitsuhashi *et al*., 2020). Furthermore, we found that longer DWI-based streamline length was associated with delayed CCSR latency and smaller amplitude measured in the contralateral hemisphere (**Fig. 6**). The findings mentioned above are in line with our recent study of intrahemispheric effective connectivity (Silverstein *et al*., 2020). A study of healthy individuals using posterior-tibial somatosensory-evoked potential likewise reported that the peak latency was correlated with the height of participants (Chu, 1986). Our sEEG sampling was restricted to the anterior head regions because the Phase-1 evaluation with scalp EEG and imaging studies indicated that the SOZ should be in these areas. Thus, further validation using sEEG data sampled from large cohorts of patients are encouraged. Large datasets may provide us with the opportunity to determine the interhemispheric connectivity across the posterior head regions.

We utilized open-source DWI data collected from healthy participants as a part of the human connectome project (Yeh *et al*., 2018) since not all of our patients underwent DWI before the sEEG placement. Thus, we were unable to assess the relationship between the fractional anisotropy (a surrogate measure of myelination) and the velocity of neural propagation at an individual level. Since our patients were 12-16 years old during sEEG recording, we cannot rule out the possibility that the streamline trajectory and fractional anisotropy could have been different (Gross *et al*., 2006; Focke *et al*., 2008). However, the relationships between peak CCSR latency, peak CCSR amplitude, and streamline length replicate those from our study involving DWI and CCEP data collected from the same patients (Silverstein *et al*., 2020) indicating that our method likely produced reliable results.

One may expect that the principle of 4D tractography may be applicable for the assessment of interhemispheric propagation of epileptiform discharges, but caution is needed. Previous invasive studies showed that SPES at the SOZ elicited high-frequency activities in the surrounding areas at a latency of 100 ms and after (van ’t Klooster *et al*., 2011; Nayak *et al*., 2014; Davis *et al*., 2018; Hebbink *et al*., 2020). Investigators suspect that such induced activities mimicking interictal epileptiform discharges might result from a cortico-thalamo-cortical propagation rather than cortico-cortical propagation. Taking this issue into account, the present study limited the CCSR analysis to nonepileptic sites with a peak latency of 11-50 ms.

The velocity of neural propagation in our patient population (i.e., mean: 2.41 mm/ms and standard deviation: 1.11 mm/ms) was comparable to that reported in a recent study in which the propagation velocity was defined as [streamline length] divided by [CCEP N1 peak latency] (i.e., range: 0.2-6.0 mm/ms; Trebaul *et al*., 2018). A previous study estimated the propagation velocity based on the nerve fiber diameters measured on post-mortem histology (Caminiti *et al*., 2013); this study reported that inter-hemispheric propagation to the homotopic frontal regions would take only 5.3-6.0 ms and that the propagation velocity would range 7.7 to 9.8 mm/ms in the frontal and temporal regions of the human brain. Some investigators consider early CCSR and N1 CCEP to reflect, at least in part, excitatory post-synaptic potentials after a synaptic delay (Adrian, 1936; Barth *et al*., 1989; Douglas *et al*., 1995; Iwasaki *et al*., 2010). If the present study considered the synaptic delay of 0.5-1.1 ms (Taschenberger and von Gersdorff, 2000; Feldmeyer *et al*., 2006) and defined the propagation velocity as [streamline length] divided by [early CCSR peak latency minus synaptic delay], the mean propagation velocity was computed as 2.47-2.55 mm/ms.

## Abbreviations

CCEPs: cortico-cortical evoked potentials
CCSRs: cortico-cortical spectral responses
DWI: diffusion-weighted imaging
MNI: Montreal Neurological Institute
sEEG: stereo-electroencephalography
SOZ: seizure onset zone
SPES: single-pulse electrical stimulation
4D: four-dimensional

## Acknowledgments

We are grateful to Karin Halsey, BS, REEGT. and Jamie MacDougall, RN, BSN, CPN at Children’s Hospital of Michigan for the collaboration and assistance in performing the studies described above. This work was supported by the National Institutes of Health [grant number: NS064033 (to E.A.); NS089659 (to J.W.J.)] and JST CREST [grant number: JPMJCR1784 (to T.M.)].

